# EZH2 inhibition enhances the activity of platinum chemotherapy in aggressive-variant prostate cancer

**DOI:** 10.1101/2024.09.06.611612

**Authors:** Maryam Latarani, Perla Pucci, Mark Eccleston, Massimiliano Manzo, Priyadarsini Gangadharannambiar, Ilaria Alborelli, Vera Mongiardini, Namra Mahmood, Mario Paolo Colombo, Benedetto Grimaldi, Sushila Rigas, Shusuke Akamatsu, Cheryl Hawkes, Yuzhuo Wang, Elena Jachetti, Francesco Crea

## Abstract

**Background:** EZH2 promotes aggressive-variant prostate cancer (AVPC) progression *via* histone H3-Lysine-27 tri-methylation (H3K27me3). We hypothesize that epigenetic reprogramming *via* EZH2 inhibitors (EZH2i) improves the efficacy of chemotherapy in AVPC.

**Methods:** We studied the expression of EZH2 in clinical prostate cancer cohorts (bioinformatics). We determined the effect of EZH2i on both cellular- and cell-free-H3K27me3 levels. We measured effects of carboplatin with/without EZH2i on AVPC cell viability (IC_50_). We studied how EZH2i modulate gene expression (RNA Seq).

**Results:** EZH2 was significantly up-regulated in AVPC vs other prostate cancer types. EZH2i reduced both cellular and cell free-H3K27me3 levels. EZH2i significantly reduced carboplatin IC_50_. EZH2i reduced the expression of DNA repair and increased the expression of pro-apoptotic genes.

**Article Highlights:**

- Polycomb-mediated gene silencing promotes prostate cancer progression
- Aggressive-variant prostate cancers (AVPCs) are characterized by increased activity of the Polycomb-Repressive Complex 2 (PRC2)
- Here we show that PRC2 inhibitors are scarcely effective as monotherapy in ACPC cells
- However the combination of PRC2 inhibitors and carboplatin is highly synergistic
- RNA Seq studies revealed that PRC2 inhibitors enhance carboplatin activity by modulating several key pathways, including DNA repair and apoptosis.

## 1. Introduction

Prolonged hormonal therapies induce phenotypic plasticity and multi-drug resistance in advanced prostate cancer (1). Malignancies emerging from this phenomenon are androgen receptor (AR)-negative or AR-indifferent and have been classified as: (I) neuroendocrine prostate cancers (NEPCs), which express typical trans-differentiation markers (e.g. synaptophysin, FOXA2 (2)); (II) double negative prostate cancers, which are both AR-negative/-indifferent and NEPC-negative (3); (III) aggressive-variant prostate cancers (AVPCs), which include adenocarcinomas and NEPCs that meet clinical and molecular criteria for enhanced metastatic ability and for androgen indifference (4). Here, we will use the broader term “AVPC” to refer to all the aforementioned malignancies. Despite extensive research efforts, there is no effective treatment for AVPC. The most common palliative treatment for these patients is platinum-based therapy, which typically results in >50% objective response rates but does not significantly improve overall survival chances (the median overall survival for these patients is shorter than 24 months) (5).

The enhancer of zeste homolog 2 (*EZH2*) locus encodes a methyltransferase that acts as part of the Polycomb-repressive complex 2 (PRC2) (6). The other canonical components of PRC2 are the proteins encoded by Embryonic Ectoderm Development (*EED*) and Suppressor of zeste 12 (*SUZ12*). In the nucleus, PRC2 induces gene silencing via histone H3-Lys27 trimethylation (H3K27me3), a repressive post translational modification. EZH2 was first described as a driver of prostate cancer progression by Varambally et al (7). Among other results, this study showed that EZH2 silencing caused a dramatic reduction in AR-negative prostate cancer cell proliferation. We and others showed that EZH2 is up-regulated in NEPC and drives prostate cancer metastasis, phenotypic plasticity and proliferation (8–10). Recently, several inhibitors of EZH2 have been developed and tested in clinical trials (11). Notably, one of these inhibitors is being tested in combination with hormonal therapies in prostate adenocarcinoma (Clinical Trial: NCT03480646). However, the role of EZH2 inhibitors in AVPC therapy, particularly in combination with platinum agents, has never been explored. Here we used prostate cancer cell lines representing the main sub-types of AVPC, including DU-145 which is AR- and NEPC-negative (12) and OPT7714 which is AR-indifferent and expresses typical NEPC markers (13). Given the heterogeneity of AVPC, it is important to test new therapies in combination with diagnostic technologies that can facilitate patient selection and treatment monitoring. We have recently demonstrated the potential clinical usefulness of a quantitative immunoassay technology that measures nucleosomal H3K27me3 levels in biological fluids (including plasma) (14). Here we have tested this technique to measure nucleosomal H3K27me3 levels in cell supernatants and compared it with the traditional methodology of measuring H3K27me3 in cellular extracts by western blotting. This approach has allowed us to determine whether the nucleosome immunoassay technology can accurately predict the pharmacodynamic activity of EZH2 inhibitors.

Our results indicate that EZH2 inhibitors potentiate the anticancer activity of platinum agents and provide a potential mechanistic framework to explain this observation.

## 2. Materials and Methods

### Gene Expression Datasets

Bioinformatic analyses were conducted through the CbioPortal dataset (<u>cBioPortal for Cancer Genomics</u>). The correlation between PRC2 gene expression and AR-activity/NEPC features e was investigated in the “Metastatic Prostate Adenocarcinoma-SU2C-PCF” dataset (containing both adenocarcinomas and NEPC samples). The EZH2 pathway analysis and the correlation between PRC2 genes and NEPC markers was performed by analysing the Cbioportal “Neuroendocrine Prostate Cancer-Multi Institute” dataset.

### Cell culture

The human cell line DU-145 (Cellosaurus Accession: CVCL_0105) is derived from a central nervous system metastasis and is a model of double-negative prostate cancer since it is both AR-negative and NEPC marker-negative (American Type Culture Collection (ATCC)). Being AR-negative and derived from a soft tissue metastasis, DU-145 cells are a clinically relevant model of AVPC (4). DU-145 cells were cultured in RPMI 1640 (Gibco®, Thermo Fisher) supplemented with 10% (v/v) of heat-inactivated Foetal Bovine Serum (FBS) (Thermo Fisher) and 1% of Penicillin-Streptomycin (Pen/Strep) (Gibco®, Thermo Fisher).

AR-negative, OPT7714 (murine cell line) is a model of NEPC generated at Istituto *Nazionale dei Tumori* (13). These cells were cultured in DMEM, high Glucose (Gibco®, Thermo Fisher) supplemented with 10% (v/v) of heat inactivated FBS (Thermo Fisher), 1% (v/v) of Pen/Strep (Gibco®, Thermo Fisher), sodium pyruvate (Gibco®, Thermo Fisher), and HBSS. No calcium, magnesium, nor phenol red (Gibco®, Thermo Fisher) was used in the culture medium.

Results obtained with the two aforementioned cell lines were further validated in human Kucap-13 cells (Cellosaurus accession CVCL_C0UV), which are a newly characterised patient-derived NEPC cell line. This cell line is the first human model of treatment-emergent NEPC. Kucap-13 cells were cultured as described in the original publication (15).

### EZH2 inhibitors

Three clinically tested EZH2 inhibitors were purchased from MedChemExpress: Tazemetostat (HY-13803), GSK-126 (HY-13470), and CPI-1205 (HY-100021).

### Cell quantification

Cells were trypsinised and collected for cell count by Luna automated cell counter (Logos Biosystems) in accordance with the manufacturer’s instructions.

### H3K27me3 measurement in cell extracts

Subconfluent cultured cells were treated with DMSO (vehicle), Tazemetostat (1, 5, and 10µM), GSK-126 (1, 5, and 10µM), and CPI-1205 (1, 5, and 10µM) for 72 hours. Cells were collected by scraping, lysed in RIPA lysis buffer [20 mM Tris-HCl (pH 8.0), 150 mM NaCl, 1 mM EDTA, 0.1% SDS, 1% Igepal, 50 mM NaF, 1 mM NaVO3] containing a protease inhibitor cocktail (Merck). Protein concentration of the cell lysate was measured using the Pierce™ Gold BCA protein assay kit (Thermo Fisher).

### Western blotting

The extracted proteins were resolved by gel electrophoresis on reducing SDS-polyacrylamide gels on 10–20% Tris-Glycine gels(Thermo Fisher Scientific) and transferred to nitrocellulose membranes (Fisher Scientific) following reported protocol (16). The membranes were incubated overnight at 4°C with either anti-H3K27me3 (1:1000, Cell Signalling Technology, Tri-Methyl-Histone H3 (Lys27) (C36B11) Rabbit mAb) diluted in 5% (w/v) BSA, or anti-GAPDH (1:10000, Sigma Aldrich) diluted in 5% dried milk.

### Nucleosomal H3K27me3 measurement in cell supernatant

Cell lines were exposed to Tazemetostat, GSK-126, and CPI-1205 at 10µM 24 hours after cell seeding. The supernatants were collected on day 3, 5, 7, and 10 after treatment, and the levels of nucleosomal H3K27me3 quantified by Nu.Q-H3K27me3 prototype colorimetric ELISA kit, (Belgium Volition SPRL). Nucleosomal vH3K27me3 levels were determined following the manufacturer’s instructions.

### Monotherapy with EZH2 inhibitors

Cell viability was determined for each cell line following exposure to different concentrations (25, 10, 1, and 0.01 µM) of EZH2 inhibitor or vehicle control (DMSO) for 5 days. The medium was replaced with fresh medium the same EZH2 inhibitors for another 5 days (the treatment lasted a total of 10 days).

### Combination of chemotherapy agent and EZH2 inhibitor

To estimate the effect of EZH2 inhibition in combination with Carboplatin (Sigma), we selected 10 µM GSK-126 because this concentration resulted in almost complete H3K27me3 ablation in at least one cell line, whilst not causing significant cell growth inhibition. Before testing the combination, we pre-treated cells with the EZH2 inhibitor for three days to ensure the occurrence of epigenetic reprogramming.

Hence, each cell line was treated with GSK-126 at 10 µM for 72 hours. Subsequently, the GSK-126 containing supernatant was replaced with a combination of GSK-126 (10µM) and Carboplatin (500, 100, 50, 10, 1, 0.1, 0.01 µM) for another 72 hours. The IC_50_ for each cell line was calculated using trypan blue exclusion cell counts. Caspase 3/7 activity was measured using the Promega Caspase-Glo assay and following Manufacturer’s instructions.

### RNA extraction and RNA sequencing

Total RNA from DU-145 cells treated with GSK-126 or vehicle was extracted using the RNeasy Mini Kit in accordance with the manufacturer’s instructions (Qiagen). NGS libraries were prepared with the Ion AmpliSeq Library Kit Plus (Thermo Fisher Scientific). For each sample, 100 ng of total RNA was reverse transcribed using the SuperScript VILO cDNA synthesis Kit (Thermo Fisher Scientific). The resulting cDNA was then amplified with the AmpliSeq Human Transcriptome Gene Expression Kit panel, targeting over 20,000 genes (> 95% of the RefSeq gene database). Amplicons were digested with the proprietary FuPa enzyme (Thermo Fisher Scientific) to generate compatible ends for barcoded adapter ligation. The resulting libraries were purified using AmpureXP beads (Agencourt) at a bead to sample ratio of 1.5X and eluted in 50 µL low TE buffer. Libraries were then diluted 1:10000 and quantified by qPCR using the Ion Universal Quantitation Kit (Thermo Fisher Scientific). Individual libraries were diluted to a concentration of 50 pM, combined in batches of 8 libraries, loaded on an Ion 540™ chip using the Ion ChefTM instrument and sequenced on an Ion S5™ (Thermo Fisher Scientific). Raw data was processed automatically on the Torrent ServerTM and aligned to the reference human genome reference build hg19 using the hg19 AmpliSeq Transcriptome fasta reference. Quality control was manually performed for each sample based on the following metrics; number of reads per sample >14X106, valid reads > 95%. RNA Seq data were uploaded in the SRA database (NCBI): PRJNA1036550.

### RNA-Seq analysis

Differential gene expression was performed using DESeq2 (17) and log-fold changes were shrunk using apeglm (18).Gene Set Enrichment Analysis (GSEA) was performed using clusterProfiler (19). Gene-pathway information was taken from the Molecular Signatures Database (MSigDB) package. Pathways with adjusted p value (p.adjust) values below 0.05 were retained for downstream analysis and visualisation.

### Statistical analyses

Experiments were conducted in three biological replicates. Data were assessed for normality using the Shapiro-Wilk test. Expression of mRNA levels from CbioPortal data and IC_50_ values were analysed using a two-tailed Student’s t-test. H3K27me3 levels were analysed using either a one-way ANOVA with Dunnett’s post hoc test or a Kruskal-Wallis with Dunn’s post hoc test. Data represent mean ± SEM and statistical analysis was performed via Graph Pad Prism 8.

## 3. Results

### 3.1. Clinical significance of PRC2 genes in aggressive-variant prostate cancer

To investigate the clinical significance of the three main components of PRC2 (EZH2, EED, SUZ12), we queried a public dataset containing histopathological, clinical, and transcriptomic information on a cohort of metastatic prostate cancer patients (20). Since the two main types of AVPC are characterised by reduced AR activity and/or NEPC histopathological features, we analysed the association between each PRC2 gene and these two characteristics. All three PRC2 genes were significantly upregulated in clinical samples with histopathological features of NEPC, compared to samples with no NEPC features (Figure 1A). We also found a significant negative correlation between the expression of *EED* and *EZH2* (but not *SUZ12*) and the “AR activity score” of clinical samples (Figure 1 B-D). This score is a transcriptome-based estimation of AR activity, which has been calculated as previously described (21).

**Figure 1.**
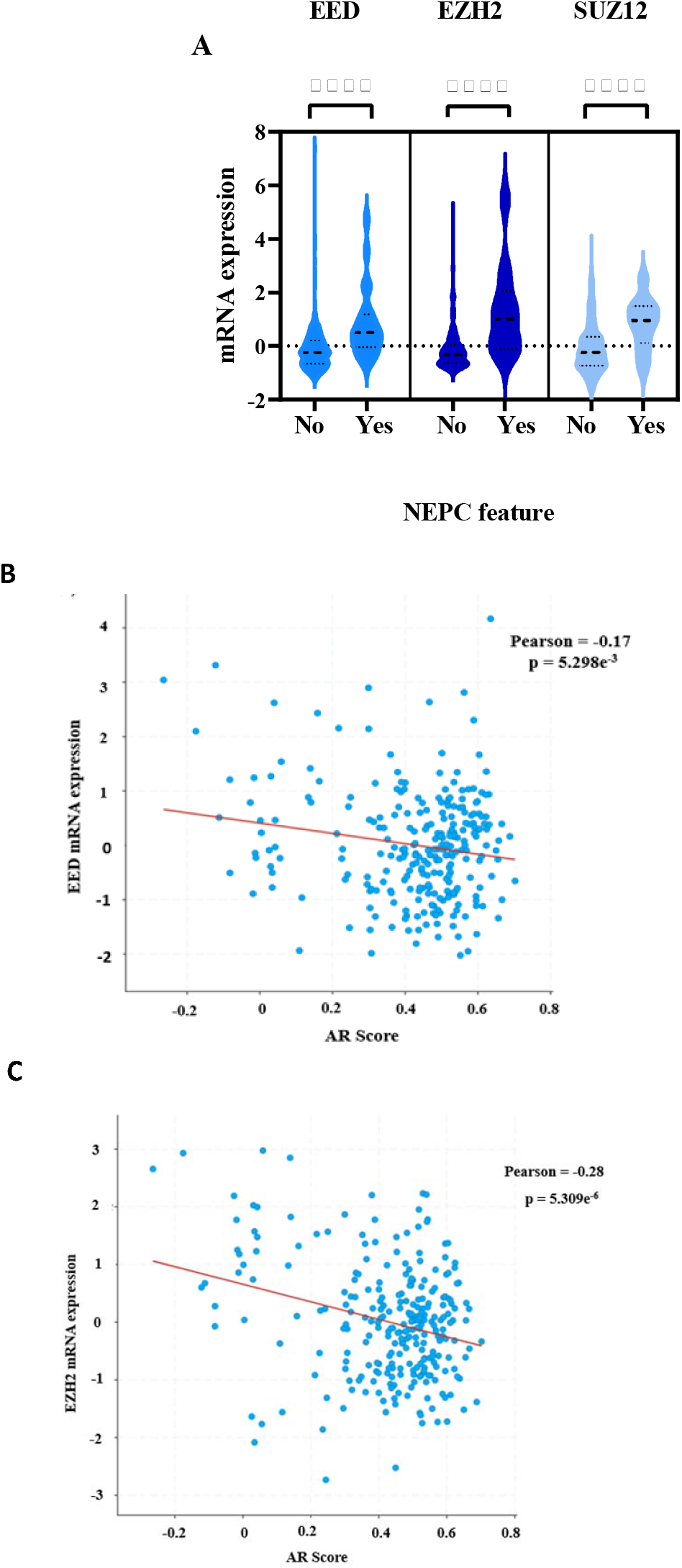
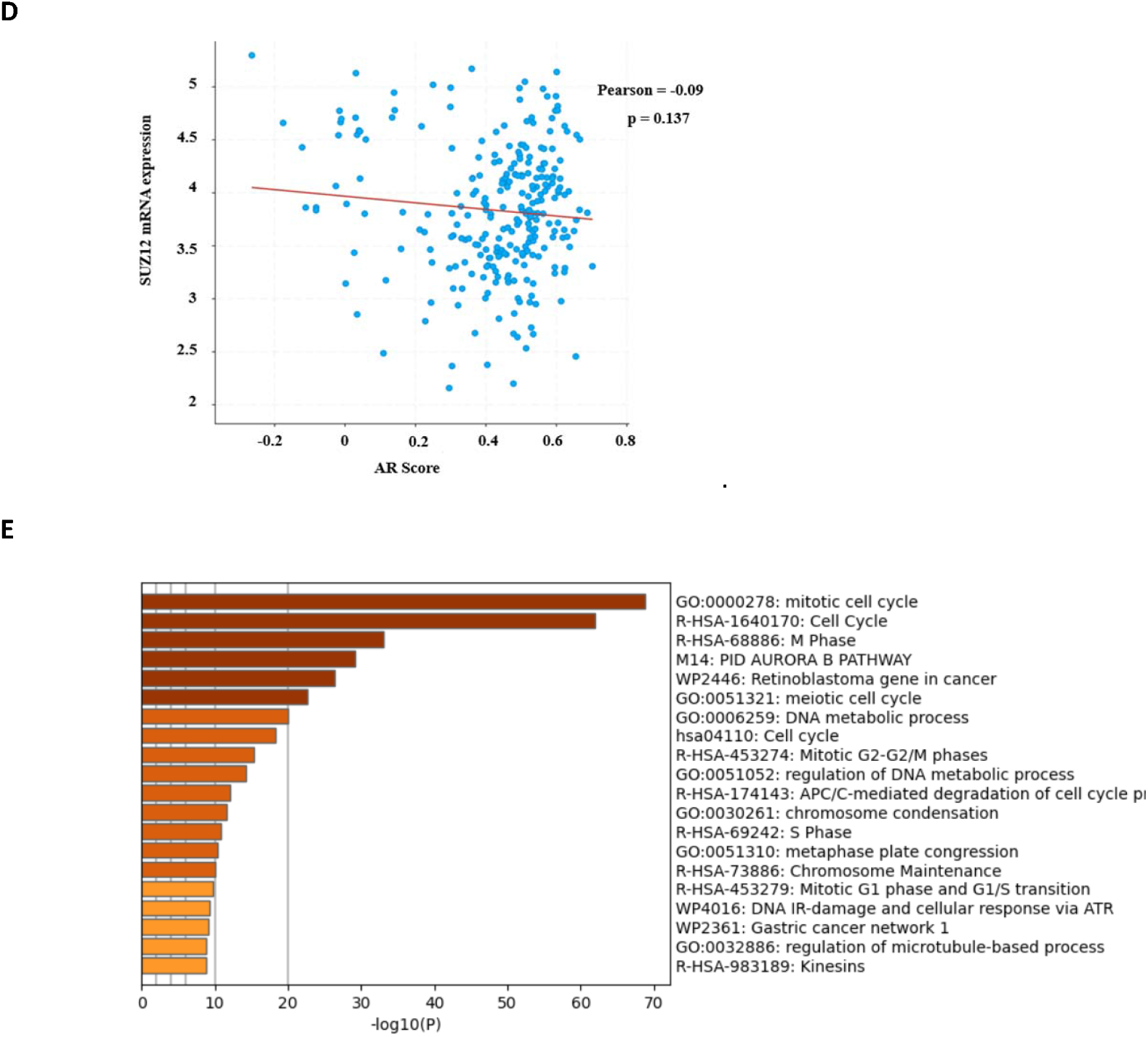
**(A)** Expression of *EED, EZH2*, and *SUZ12* mRNA levels in n=264 PCa samples (CbioPortal), <u>Neuroendocrine Prostate Cancer-Multi Institute dataset</u> accessed on 10/01/2023) with or without NEPC features. **** = p <0.0001, two-tailed Student t-test,. Correlation between gene expression and AR activity score for genes (**B**) EED, (**C**) EZH2 and (**D**) SUZ12. Analises for a-d were conducted using the <u>Metastatic Castration Resistant Prostate Cancer dataset</u> on Cbioportal (**E**). Pathway analysis of genes significantly co-expressed with EZH2 in the Cbio-portal <u>Neuroendocrine Prostate Cancer-Multi Institute dataset</u> (cases were defined based on cell morphology and expression of NEPC makers). Genes with Pearson correlation coefficient>0.85 (EZH2) were downloaded in Metascape for pathway analysis (https://metascape.org/gp/index.html#/main/step1).

We then conducted a transcriptome analysis on a different dataset of clinical NEPC samples, to identify pathways that were commonly associated with EZH2 expression. This analysis showed that some key oncogenic pathways were significantly associated with EZH2 expression in NEPC samples (Figure 1E). Most of the enriched pathways were correlated with cell cycle (G2 and M phase), DNA repair, and chromosome maintenance. The same dataset was interrogated to study correlations between PRC2 gene expression and individual NEPC markers (*SYP, ENO2*). The analysis provided an independent confirmation of the strong positive correlation between PRC2 genes and almost NEPC markers (5 out of 6 significant correlations, Tab. 1).

**Tab 1.** Linear correlation results from the Neuroendocrine Prostate Cancer (Multi-Institute, Nat Med 2016, https://www.cbioportal.org/study/summary?id=nepc_wcm_2016) Cbio portal study. We have computed linear correlation analyses between the mRNA expression of PRC2 genes and the mRNA expression two typical NEPC markers: synaptophysin (*SYP*) and neural specific enolase (*ENO2*). Q values are derived from the Benjamini-Hochberg FDR correction procedure.

| GENES | SPEARMAN'S CORRELATION | Q VALUE |
| --- | --- | --- |
| *EZH2-ENO2* | 0.744 | 5.92e<sup>-8</sup> |
| *EZH2-SYP* | 0.578 | 1.427e<sup>-4</sup> |
| *EED-ENO2* | 0.719 | 5.07e<sup>-8</sup> |
| *EED-SYP* | 0.544 | 1.720e<sup>-4</sup> |
| *SUZ12-ENO2* | 0.482 | 2.183e<sup>-3</sup> |
| *SUZ12-SYP* | 0.178 | 0.310 |

### 3.2. EZH2 inhibitors are effective in AVPC cells

In light of the bioinformatic results, we investigated whether the three clinically tested EZH2 inhibitors Tazemetostat, GSK-126, and CPI-1205 were active in NEPC cells (OPT7714) and in AR-negative cells that do not express NEPC markers (DU-145) (22). Western blot results demonstrated that all EZH2 inhibitors reduced H3K27me3 (a marker of PRC2 activity), although to a variable degree between cell types. In DU-145 cells, GSK-126 significantly decreased H3K27me3 compared to cells exposed to vehicle (Figure 2A), whilst an almost complete suppression of H3K27me3 expression was observed in OPT7714 cells at concentrations of 5 µM or higher of Tazemetostat, GSK-126 and CPI-1205 (Figure 2B). We also tested whether the reduction in H3K27me3 was due to a reduction in total histone H3 levels. The ratio of Nucleosomal H3K27me3/total nucleosomal H3.1 was determined in cells treated with GSK-126 or DMSO control. Our results show that the relative levels of H3K27me3/total H3.1 were significantly reduced in cells exposed to the EZH2 inhibitor (Suppl. Fig.1).

**Figure 2.**
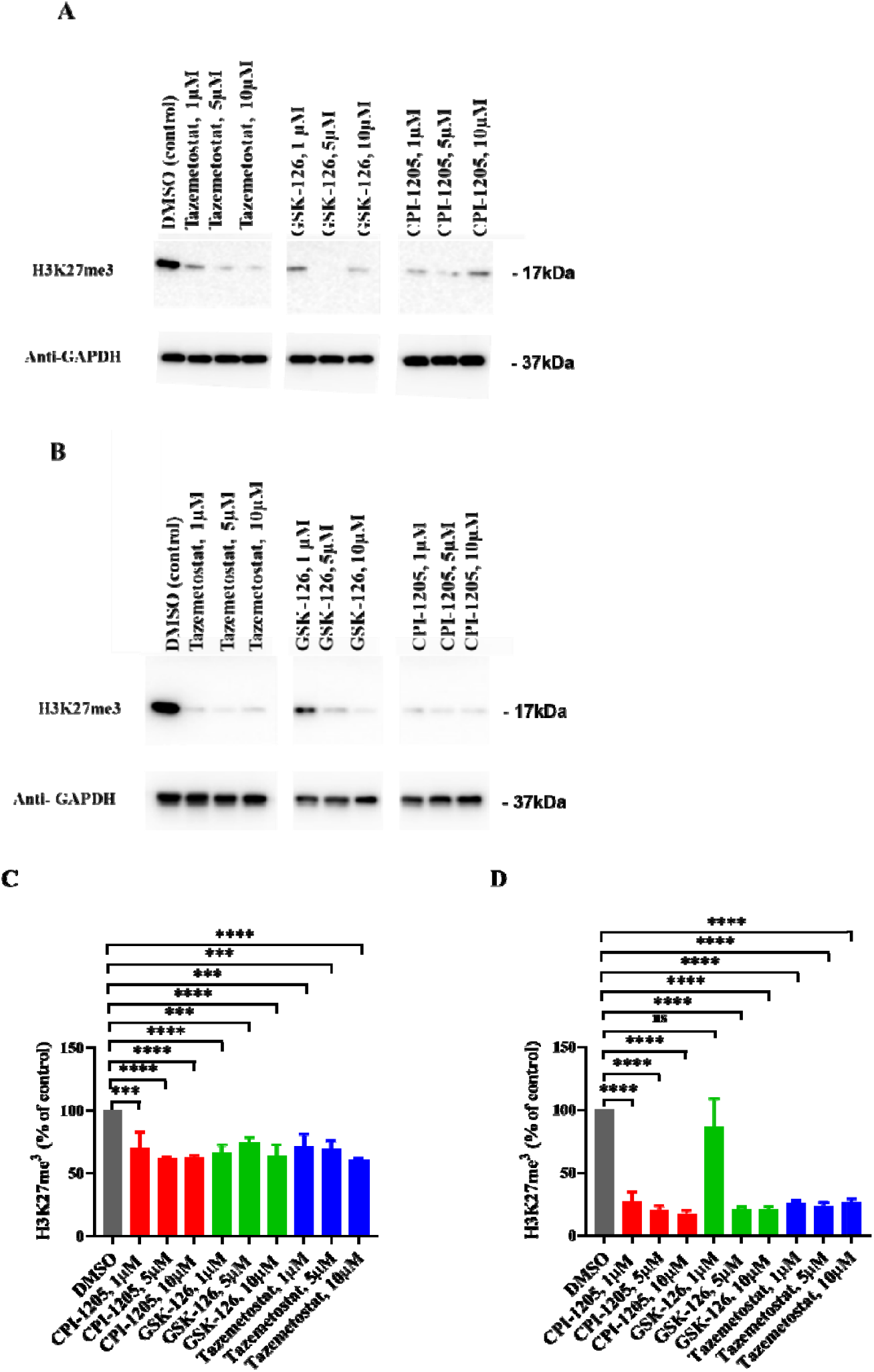
H3K27me3 levels in DU145 (**A** and **C**) and OPT7714 cells (**B** and **D**) exposed to different EZH2 inhibitors (upper panel representative image; lower panel statistical analysis of three quantified blots). Cells were treated with DMSO (vehicle), Tazemetostat 1, 5, and 10µM, GSK-126 1, 5, and 10µM, and CPI-1205 1, 5, and 10µM for 72 hours; cellular protein extracts were analysed by immunoblotting using anti-H3K27me3 and anti-GAPDH antibodies. Nu.Q Volition-measured H3K27me3 levels in the supernatant of DU-145 (C) and OPT7714 (D) cells in the same conditions described above. ***P< 0.001 and ****P< 0.0001 one-way ANOVA with Dunnett post-hoc test (C and D).

To explore whether levels of nucleosomal H3K27me3 in biological fluids could predict the activity of the EZH2 inhibitors, we used a novel ELISA (Nu.Q) to measure the levels of this histone modifications in cell-free nucleosomes from cellular supernatants. Treatment of AVPC cells with the EZH2 inhibitors caused a significant reduction in cell-free nucleosomal H3K27me3 levels (Figure 2C, D). Notably, the reduction in cell-free H3K27me3 levels was more pronounced in OPT7714 than in DU-145 cells, which was consistent with the Western blot results from cellular extracts.

### 3.3. EZH2 inhibition increases the anticancer activity of carboplatin in AVPC cells

Having confirmed the pharmacodynamic activity of the three EZH2 inhibitors on H3K27me3 levels, we then assessed their anti-proliferative effects. As shown in Figure 3A and B, treatment with EZH2 inhibitors alone (without chemotherapy) reduced the cell count of AVPC cells only at higher concentrations, particularly with GSK-126. We then exposed the cells to a fixed dose of GSK-126 (10 µM), followed by carboplatin at different concentrations, or carboplatin alone (Figure 3C and D). As shown in Figure 4E and F, the combination treatment showed a significant IC_50_ reduction in both cell lines, compared to carboplatin monotherapy.

**Figure 3.**
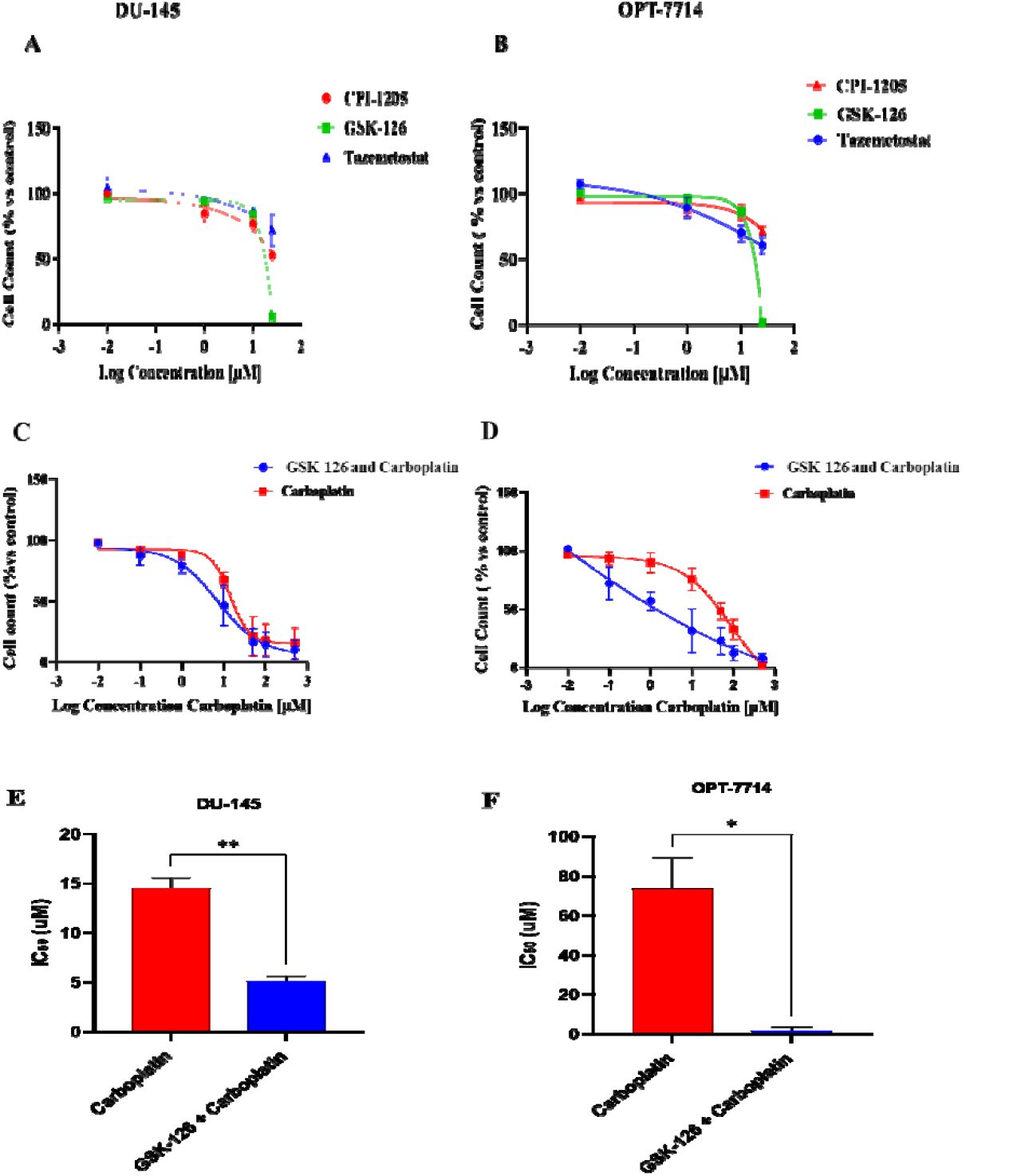
Growth inhibitory effects of Tazemetostat, GSK-126, and CPI-1205 on DU-145 (**A**) and OPT7714 (**B**) cells. The cells were counted after exposure to different concentrations of EZH2 inhibitor (0.01, 1, 10, and 25 µM), or DMSO for 10 days. Cell viability of DU-145 (**C**) and OPT7714 (**D**) treated with combination of GSK-126 and Carboplatin vs Carboplatin alone. Cells were exposed to 10 µM GSK-126 for 72 hours, followed by GSK-126 (10 µM) and Carboplatin (different concentrations as indicated by the X axis) for the following 72 hours. Cells were counted at the end of the experiment. IC_50_ values of DU-145 (**E**) and OPT7714 (**F**) treated with a combination of GSK-126 and Carboplatin vs Carboplatin alone. *P<0.05, **P<0.01, two-tailed Student’s t-test.

**Figure 4.**
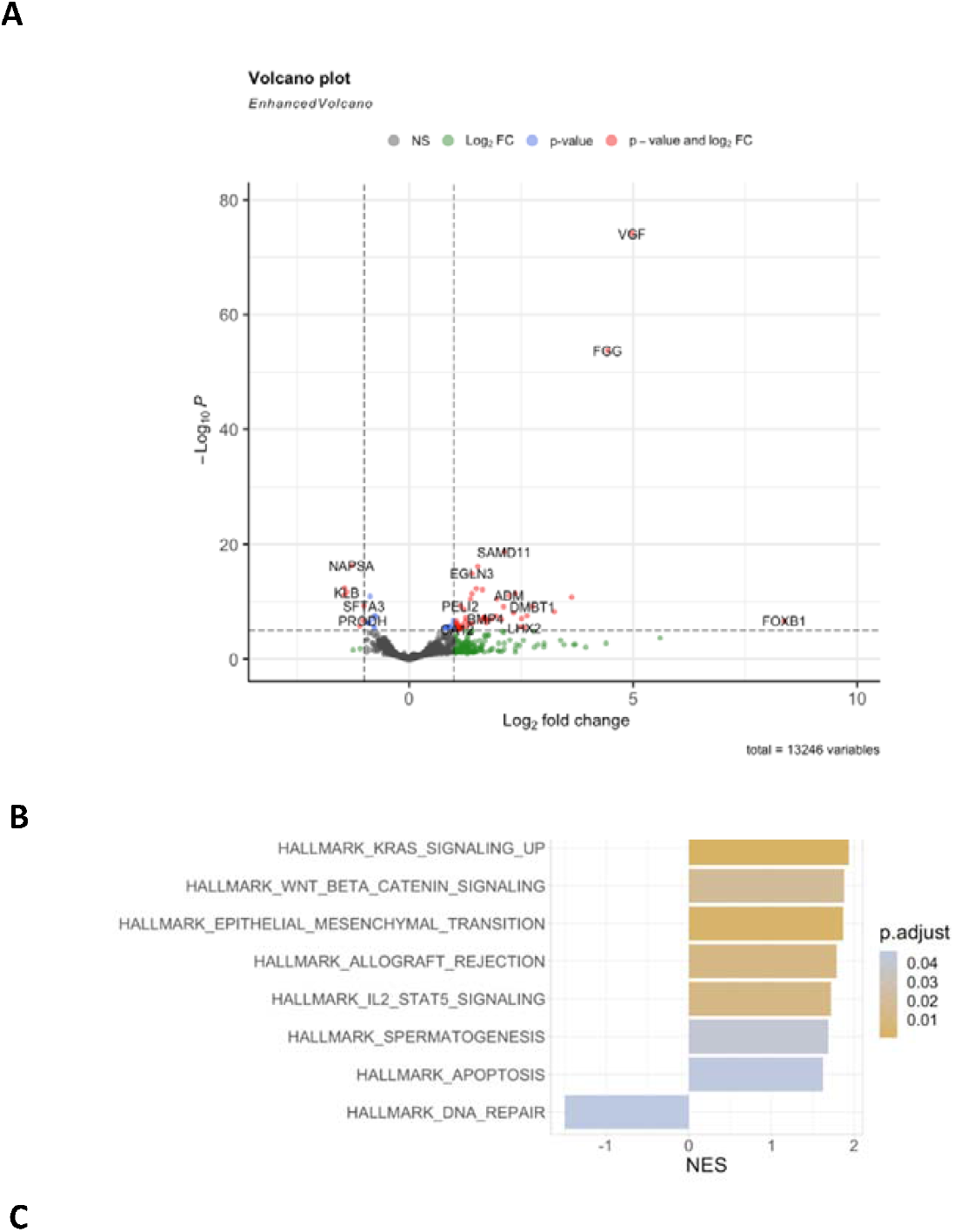
Transcriptome analysis of DU-145 cells exposed to EZH2 inhibitor. (**A**) Volcano plot of RNA Seq from cells exposed to GSK-126 vs control. Differentially expressed genes are coloured in red if they are above the fold-change and -Log (p.adjust-value) threshold; genes coloured in green are only above the -Log (p.adjust value) threshold. The names of some highly up- and down-regulated genes are shown in the plot. (**B**) A comparison of normalised enrichment scores (NES) that are significantly different in cells exposed to GSK-126 or control. A positive NES indicates up-regulation of the pathway in cells treated with GSK-126. A negative NES indicates down-regulation of the pathway in cells treated with GSK-126.

**Figure 5.**
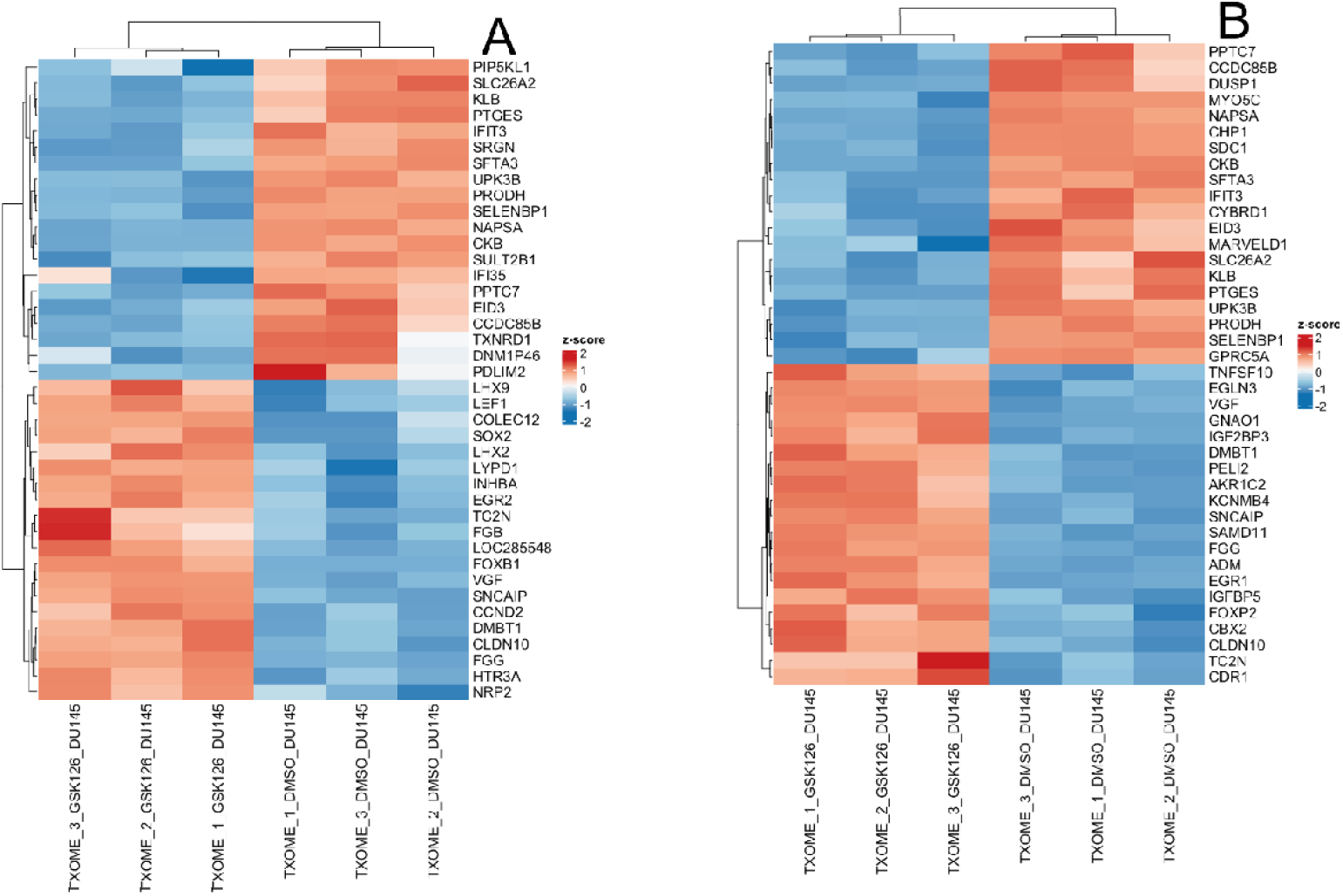
Heatmap from DU-145 RNA Sequencing results. Genes were ranked based on decreasing fold-change (A) or increasing p.adjust (B) and subsequently grouped by hierarchical clustering. Red colour signifies up-regulation, blue colour down regulation. Results are shown as single values in each of the biological triplicates for treatment with GSK-126 or DMSO (control).

To further corroborate our hypothesis, we sought to replicate our findings in a patient-derived model of treamtnet emergent NEPC (Kucap-13). These cells were treated with either carboplatin alone or carboplatin plus GSK-126 at a fixed dose. We found that also in this model (which grows in spheroid-like aggregates) the combination induced a significant reduction in cell viability, compared to carboplatin only (Suppl. Fig. 2). Notably, EZH2 inhibitor monotherapy does not show significant anti-proliferative effects in this cell line (15). Taken together, these results show that whilst EZH2 inhibitors might have limited effects as monotherapy, they significantly enhance the anti-proliferative activiyty of carboplatin in AVPC cells.

### 3.4. EZH2 inhibition reduces the activity of DNA repair genes

To provide a molecular explanation for the cellular response shown in Figure 3, we performed RNA sequencing on DU-145 cells exposed to GSK-126 vs control (DMSO). This analysis confirmed that EZH2 inhibition causes a profound alteration of the cell’s transcriptome. Among the differentially expressed genes (DEGs), we observed more up-regulated than down-regulated transcripts upon EZH2 inhibition (Figure 4A). However, DEGs contained a mix of oncogenic and onco-suppressive genes. A GSEA revealed that EZH2 inhibition modulates several biological functions: in most cases this led to the activation of pathways, as shown by positive enrichment scores (Figure 4B). The balance between the activation of pro-apoptotic and anti-apoptotic (e.g., *KRAS*) pathways provides a potential explanation for the modest effects of EZH2 inhibitor monotherapy on cell growth. Notably, we found that EZH2 inhibition significantly reduced the activity of DNA repair pathways (Figure 4B; negative enrichment score). Amongst the down-regulated DNA repair genes in this pathway we have identified *ERCC1, ERCC2* and *P53*.

It is therefore conceivable that the GSK-126-dependent epigenetic reprogramming modulates several drug-sensitising pathways, leading to increase sensitivity to carboplatin.

## 4. Discussion

In this study, we have shown that EZH2 inhibition can increase AVPC cell sensitivity to carboplatin. Our experiments have been carried out in cell lines that represent different subtypes of AVPC (neuroendocrine and double negative) and could therefore represent a substantial fraction of clinical metastatic prostate cancers. The combination of GSK-126 and carboplatin effectively reduced live cell count compared to the agents alone in three different preclinical models, showing at least additive effects. The translational potential of this combination should be addressed in further preclinical studies, where the pharmacokinetics of the treatments should be tested in preclinical models and patient-derived models.

EZH2 inhibitors potentiate the activity of several anticancer drugs. For example, we showed that the first EZH2 inhibitor (DZNeP) increases pancreatic cancer sensitivity to gemcitabine (23). More recently, the combination between GSK-126 and Ataxia-Telangiectasia Mutated (ATM) inhibitors was identified as synthetically lethal in breast cancer (24). PRC2 directly silences several anticancer genes, whilst also indirectly activating numerous cancer-promoting pathways (25). This paradigm is reflected by our RNA Seq data, showing that EZH2 inhibition resulted in both gene reactivation (most likely because of reduced H3K27me3 at specific *loci*) and gene down-regulation (most likely as an indirect or non-canonical effect). It is therefore conceivable that EZH2 inhibitors can have diverse effects on different cancer cell types. In this study, we showed that EZH2 inhibition impairs the expression of several DNA repair genes that are responsible for resistance to platinum-based chemotherapy. For example, ERCC1 is directly involved in the nucleotide excision repair pathway, which is activated by platinum-dependent DNA damage (26). Intra-tumoral ERCC1 expression is inversely correlated with clinical outcomes in small cell lung cancer patients treated with cisplatin, where lower expression is correlated with better response (27). Similarly, TP53 inactivation has been linked to platinum sensitivity (28). This could be due to ineffective DNA damage checkpoint, which leads to cell death upon platinum treatment (29). Notably, additional analysis of our RNA-Seq data indicated that activation of pro-apoptotic pathways could synergize with reduced DNA repair, thereby providing a further explanatory framework for the observed interaction between EZH2 inhibitors and carboplatin.

Taken together, these results suggest that EZH2 inhibition, whilst not lethal *per se*, induces significant vulnerabilities in AVPC cells. It would be interesting to systematically explore other synthetically lethal combinations in this incurable cancer.

Our results also suggest a significant positive correlation between cell-free and cellular H3K27me3 post translational modification levels in nucleosomes. We described the potential prognostic value of circulating nucleosomes (14). Here, we show that cell-free nucleosomal H3K27me3 can be detected in the supernatant from AVPC cell culture and that H3K27me3 levels decrease upon treatment with EZH2 inhibitors. Notably, the levels of cell-free H3K27me3 nucleosomes measured with the Nu.Q assays are consistent with the results obtained with standard western blot from cellular protein extracts. This suggests that the nucleosome immunoassay technology could be useful for patient selection and therapeutic monitoring. We have proposed this strategy in a previous publication (30).

Our *in vitro* evidence shows that EZH2 inhibition enhances the activity of platinum-based chemotherapies; however, further studies are needed to clarify how EZH2 inhibitors affect the local chromatin status of specific genetic *loci*, and how this translates into transcriptomic changes leading to reduced DNA repair efficiency. Our results also indicate that PRC2 genes are over-expressed in both NEPC and AR-negative cancers. However, these transcriptomic data (Figure 1) will need to be confirmed at protein level to increase the translational significance of our findings.

Another potential limitation of our study relates to the use of clinically available EZH2 inhibitors. We have decided to use these compounds because we wanted to generate clinically relevant results. In particular, we have selected three compounds that were being used in clinical trials at the beginning of our study. However, currently available EZH2 inhibitors have some limitations. It is well known that EZH2 has three main functions: the canonical PRC2-dependent H3K27 methyl-transferase activity; non-canonical methyl-transferase activities (which are PRC-2-independent and target non-histone proteins); methylation-independent functions such as interacting with transcription factors and signaling molecules (31). Whilst all these functions may be relevant to prostate cancer progression, clinically tested EZH2 inhibitors block the canonical and non-canonical methyl-transferase activity of EZH2, but non the methylation-independent functions (11). Hence it is conceivable that future inhibitors, with a broader spectrum of action will be more effective in AVPC.

Further research could also elucidate the role of EZH2 inhibitors alone or in combination with chemotherapy *in vivo* to increase the translational clinical potential of this study.

## 5. Conclusions

These results suggest that clinically tested EZH2 inhibitors increase carboplatin sensitivity in AVPC cells and that combinations of these compounds may be effective in different subtypes of AVPC. EZH2 inhibition, using three different drugs, strongly reduces H3K27me3 levels in AVPC cells and may affect DNA repair and other cancer-relevant pathways. We also showed that the activity of EZH2 inhibitors could be monitored by a novel nucleosome immunoassay technology, via detection of secreted levels of H3K27me3-modified nucleosomes from AVPC cells.

## Author Contributions

ML: investigation, formal analysis; PP: investigation, formal analysis, writing-first draft and editing; ME: conceptualization, writing-editing; MM and IA: data curation and formal analysis, writing-editing; NM: formal analysis, writing-editing; PG: validation and formal analysis; MPC: methodology, writing-editing; VM: validation and formal analysis; BG: validation and supervision, writing-editing; SR and CH: supervision, writing-editing; SA: methodology, writing-editing; YW: conceptualization, supervision; EJ: methodology, writing-editing; FC: supervision, conceptualization, writing-first draft.

## Data Availability Statement

RNA Seq data were uploaded in the SRA database (NCBI): PRJNA1036550.

## References

1. Beltran H, Demichelis F. Therapy considerations in neuroendocrine prostate cancer: What next? Endocr Relat Cancer. 2021;28:T67–78.

2. Mather RL, Parolia A, Carson SE, Venalainen E, Roig-Carles D, Jaber M, et al. The evolutionarily conserved long non-coding RNA LINC00261 drives neuroendocrine prostate cancer proliferation and metastasis via distinct nuclear and cytoplasmic mechanisms. Mol Oncol. 2021;15:1921–41.

3. Brown LC, Halabi S, Somarelli JA, Humeniuk M, Wu Y, Oyekunle T, et al. A phase 2 trial of avelumab in men with aggressive-variant or neuroendocrine prostate cancer. Prostate Cancer Prostatic Dis. 2022;25:762–9.

4. Spetsieris N, Boukovala M, Patsakis G, Alafis I, Efstathiou E. Neuroendocrine and aggressive-variant prostate cancer. Cancers (Basel). 2020;12:1–20.

5. Ruiz de Porras V, Font A, Aytes A. Chemotherapy in metastatic castration-resistant prostate cancer: Current scenario and future perspectives. Cancer Lett [Internet]. Elsevier B.V.; 2021;523:162–9. Available from: 10.1016/j.canlet.2021.08.033

6. German B, Ellis L. Polycomb Directed Cell Fate Decisions in Development and Cancer. Epigenomes. 2022;6:1–30.

7. Varambally S, Dhanasekaran SM, Zhou M, Barrette TR, Kumar-Sinha C, Sanda MG, et al. The polycomb group protein EZH2 is involved in progression of prostate cancer. Nature. 2002;419:624–9.

8. Etienne Dardenne1,11, Himisha Beltran2,3,4,11, Matteo Benelli5, Kaitlyn Gayvert6,7, Adeline Berger1, Loredana Puca4, Joanna Cyrta1,4, Andrea Sboner1,4,6,7, Zohal Noorzad1, Theresa MacDonald1, Cynthia Cheung1, Ka Shing Yuen1, Dong Gao8, Yu Chen3,8,9, Marti, Puca L, Cyrta J, Sboner A, Noorzad Z, MacDonald T, et al. N-Myc induces an EZH2-mediated transcriptional program driving Neuroendocrine Prostate Cancer. Cancer Cell. 2016;30:563–77.

9. Clermont PL, Lin D, Crea F, Wu R, Xue H, Wang Y, et al. Polycomb-mediated silencing in neuroendocrine prostate cancer. Clin Epigenetics. 2015;7:1–13.

10. Davies A, Nouruzi S, Ganguli D, Namekawa T, Thaper D, Linder S, et al. An androgen receptor switch underlies lineage infidelity in treatment-resistant prostate cancer. Nat. Cell Biol. 2021.

11. Duan R, Du W, Guo W. EZH2: A novel target for cancer treatment. J Hematol Oncol. Journal of Hematology & Oncology; 2020;13:1–12.

12. Korsen JA, Gutierrez JA, Tully KM, Cartera LM, Samuels Z V., Rudin CM, et al. Delta-like ligand 3–targeted radioimmunotherapy for neuroendocrine prostate cancer. Proc Natl Acad Sci U S A. 2022;119:1–7.

13. Mauri G, Jachetti E, Comuzzi B, Dugo M, Arioli I, Miotti S, et al. Genetic deletion of osteopontin in TRAMP mice skews prostate carcinogenesis from adenocarcinoma to aggressive human-like neuroendocrine cancers. Oncotarget. 2016;7:3905–20.

14. Salani F, Latarani M, Casadei-Gardini A, Gangadharannambiar P, Fornaro L, Vivaldi C, et al. Predictive significance of circulating histones in hepatocellular carcinoma patients treated with sorafenib. Epigenomics. 2022;14:507–17.

15. Okasho K, Mizuno K, Fukui T, Lin YY, Kamiyama Y, Sunada T, et al. Establishment and characterization of a novel treatment-related neuroendocrine prostate cancer cell line KUCaP13. Cancer Sci. 2021;112:2781– 91.

16. Kurien BT, Hal Scofield R. Western blotting: Methods and protocols. West Blotting Methods Protoc. 2015;1–509.

17. Love MI, Huber W, Anders S. Moderated estimation of fold change and dispersion for RNA-seq data with DESeq2. Genome Biol. 2014;15:1–21.

18. Zhu A, Ibrahim JG, Love MI. Heavy-Tailed prior distributions for sequence count data: Removing the noise and preserving large differences. Bioinformatics. 2019;35:2084–92.

19. Wu T, Hu E, Xu S, Chen M, Guo P, Dai Z, et al. clusterProfiler 4.0: A universal enrichment tool for interpreting omics data. Innovation [Internet]. Elsevier; 2021;2:100141. Available from: 10.1016/j.xinn.2021.100141

20. Abida W, Cyrta J, Heller G, Prandi D, Armenia J, Coleman I, et al. Genomic correlates of clinical outcome in advanced prostate cancer. Proc Natl Acad Sci U S A. 2019;166:11428–36.

21. Hieronymus H, Lamb J, Ross KN, Peng XP, Clement C, Rodina A, et al. Gene expression signature-based chemical genomic prediction identifies a novel class of HSP90 pathway modulators. Cancer Cell. 2006;10:321–30.

22. Fraser JA, Sutton JE, Tazayoni S, Bruce I, Poole A V. hASH1 nuclear localization persists in neuroendocrine transdifferentiated prostate cancer cells, even upon reintroduction of androgen. Sci Rep. 2019;9:1–15.

23. Avan A, Crea F, Paolicchi E, Funel N, Galvani E, Marquez VE, et al. Molecular mechanisms involved in the synergistic interaction of the EZH2 inhibitor 3-deazaneplanocin A (DZNeP) with gemcitabine in pancreatic cancer cells. Mol Cancer Ther. 2012;11:1735–1746.

24. Ratz L, Brambillasca C, Bartke L, Huetzen MA, Goergens J, Leidecker O, et al. Combined inhibition of EZH2 and ATM is synthetic lethal in BRCA1-deficient breast cancer. Breast Cancer Res [Internet]. BioMed Central; 2022;24:1–19. Available from: 10.1186/s13058-022-01534-y

25. Crea F, Fornaro L, Bocci G, Sun L, Farrar WL, Falcone A, et al. EZH2 inhibition: Targeting the crossroad of tumor invasion and angiogenesis. Cancer Metastasis Rev. 2012;31:753–61.

26. Ciniero G, Elmenoufy AH, Gentile F, Weinfeld M, Deriu MA, West FG, et al. Enhancing the activity of platinum-based drugs by improved inhibitors of ERCC1–XPF-mediated DNA repair. Cancer Chemother Pharmacol [Internet]. Springer Berlin Heidelberg; 2021;87:259–67. Available from: 10.1007/s00280-020-04213-x

27. Xie K, Ni X, Lv S, Zhou G, He H. Synergistic effects of olaparib combined with ERCC1 on the sensitivity of cisplatin in non⍰small cell lung cancer. Oncol Lett. 2021;21:1–8.

28. Lin S, Li X, Lin M, Yue WX. Meta-analysis of P53 expression and sensitivity to platinum-based chemotherapy in patients with non-small cell lung cancer. Med (United States). 2021;100:E24194.

29. Li S, Wang L, Wang Y, Zhang C, Hong Z, Han Z. The synthetic lethality of targeting cell cycle checkpoints and PARPs in cancer treatment [Internet]. J. Hematol. Oncol. BioMed Central; 2022. Available from: 10.1186/s13045-022-01360-x

30. Kang N, Eccleston M, Clermont PL, Latarani M, Male DK, Wang Y, et al. EZH2 inhibition: A promising strategy to prevent cancer immune editing. Epigenomics. 2020;12:1457–76.

31. Zimmerman SM, Lin PN, Souroullas GP. Non-canonical functions of EZH2 in cancer. Front Oncol. 2023;13:1–7.

